# Structure of an atypical homodimeric actin capping protein from the malaria parasite

**DOI:** 10.1101/2020.08.16.253187

**Authors:** Ábris Ádám Bendes, Petri Kursula, Inari Kursula

## Abstract

Actin capping proteins (CPs) are essential regulators of actin dynamics in all eukaryotes. Their structure and function have been extensively characterized in higher eukaryotes but their role and mechanism of action in apicomplexan parasites remain enigmatic. Here, we present a crystal structure of a unique homodimeric CP from the rodent malaria parasite *Plasmodium berghei*. In addition, we compare homo- and heterodimeric arrangements of *P. berghei* CPs (*Pb*CPs) in solution. We complement our findings by describing the regulatory effects of *Pb*CPs on heterologous skeletal muscle α-actin as well as parasite actin. Comprehensive kinetic and steadystate measurements show atypical regulation of actin dynamics; *Pb*CPs facilitate rapid turnover of parasite actin I without affecting the apparent critical concentration. Possibly to rescue actin filament capping in life cycle stages where the CP β-subunit is downregulated, homo- and heterodimeric *Pb*CPs show redundant effects *in vitro*. However, our data suggest that homodimers may in addition influence actin kinetics by recruiting lateral actin dimers. This unusual function could arise from the absence of a β-subunit, as the asymmetric *Pb*CP homodimer lacks the structural elements essential for canonical barbed end interactions, suggesting a novel CP binding mode. These findings facilitate further studies aimed at elucidating the precise actin filament capping mechanism in *Plasmodium* and the eligibility of *Pb*CPs as drug targets against malaria.

**Significance statement:** Malaria parasites of the genus *Plasmodium* display a unique form of gliding motility, which depends on an unconventional actomyosin motor. Actin capping proteins (CPs) play a major role in regulating parasite motility. Here, we describe a unique *Plasmodium berghei* CP (*Pb*CP) system, behaving contradictory to canonical heterodimeric CPs, more suited to regulate the fast dynamics of the parasite actin system. We present the crystal structure of a distinctive homodimeric form of *Pb*CP and extensive biochemical data, describing the atypical behavior of each *Pb*CP form. The *Pb*CP homodimer displays capping even in the absence of canonical conserved structural elements, suggesting a novel actin-CP interaction mode. These distinct structural properties could provide opportunities for drug design against malaria.

## Introduction

Actin is a major element of the eukaryotic cytoskeleton and essential for a spectrum of cellular functions, including cell motility, intracellular transport, endocytosis, and cytokinesis. The versatile behavior of actin-based systems requires tight spatial and temporal control that is conveyed by a vast number of regulatory proteins (1). Actin capping proteins (CPs) are heterodimeric proteins, ubiquitously present in eukaryotes (2). CPs play a crucial role in actin dynamics by binding with high affinity to the fast-growing barbed end of filamentous actin (F-actin) in a Ca^2+^ independent manner (3), limiting protomer exchange to the pointed end. CPs are present in various cytoskeletal structures, such as lamellipodial protrusions (4), dynactin (5), and in the sarcomere, linking microfilaments to the Z-disks (6). Most vertebrates encode multiple isoforms of CP subunits with diverse functions (2). CPs are essential for human and zebrafish morphogenesis (7) and they belong to the core set of proteins needed to reconstitute actin-based motility *in vitro* (8). The average cytosolic concentration of CP in eukaryotes is in the range of 0.5-1.5 μM (9, 10), which, considering the high affinity and 1:1 stoichiometry of CP towards actin filaments, leads to a high number of constantly capped barbed ends *in vivo* (10). However, formation of free barbed ends is essential for rapid actin network assembly (11, 12), and control of CP expression levels is required for optimal actin-based cellular functions (8, 9). Thus, the mechanism of capping/uncapping is modulated by various external factors. Steric and allosteric regulators of CPs include polyphosphoinositides (PIPs), V-1/myotrophin, and CARMIL proteins, while indirect barbed end competitors include formins and ENA/VASP proteins (2).

CPs are comprised of two subunits, CPα and CPβ, each with an approximate molecular weight of 32-36 kDa (2). In most eukaryotes, the individual subunits are conserved, although the sequence identity between the subunits is typically low (2). Despite the sequence divergence, the subunits share a strikingly similar fold, resulting in a unique quaternary structure where the two subunits take up a compact arrangement with a pseudo two-fold symmetry (13). All currently available structures of heterodimeric CPs resemble the characteristic shape of a stipitate mushroom (5, 13, 14). Three N-terminal antiparallel helices form the “stalk” domain flanked by a characteristic β-stranded “globule” domain of each monomer. The “cap” is comprised of a well-ordered arrangement of two 5-stranded antiparallel β-sheets, crested by a backbone of four helices formed conjunctly by the subunits. Emerging from the cap structure are the C-terminal “tentacles” of each subunit (termed α- and β-tentacle, respectively) containing a longer flexible loop region and an amphipathic helix (13). According to the current view, CPs bind the terminal two actin protomers of a filament in a sequential mode that involves a coordinated interplay of a positively charged patch on the cap and both tentacle domains of the subunits (15, 16).

Alongside other members of the vast phylum of *Apicomplexa*, malaria parasites (*Plasmodium* spp.) use a special actomyosin motor for motility and host cell invasion (17). These parasites have two non-canonical actin isoforms and a modest set of 10-15 actin-binding proteins, lacking identifiable orthologues of the Arp2/3 complex, the majority of capping/severing/cross-linking proteins, or regulators like ENA/VASP proteins (17, 18). Contrary to the majority of eukaryotes, *Plasmodium* spp. encode only one isoform of each CP subunit (19). Metazoan CPs characterized to date nucleate polymerization, decrease elongation rate in preseeded systems, block dilution-induced depolymerization from barbed ends, and increase critical concentration to the level of the pointed end (2). Even though individual CP subunits and isoforms are expressed at different levels during different stages and cell types (20), CP subunits are largely insoluble and non-functional when expressed alone *in vitro*, indicating that they only exist as heterodimers (21). In *Plasmodium*, however, in addition to a heterodimeric CP (19, 22), a unique homodimer form of the α-subunit exists and displays capping activity of homo- and heterologous actin filaments *in vitro* (23, 24). Here, we present the first crystal structure of a homodimeric CP from the rodent malaria parasite *Plasmodium berghei* (*Pb*CPαα). The structure demonstrates critical differences compared to canonical heterodimeric CPs. We complement our structural findings with extensive biochemical characterization of *Pb*CPαα and the *P. berghei* CP heterodimer (*Pb*CPαβ), showing that the homo- and heterodimers have distinct functions that differ from the canonical CP heterodimer (CapZαβ) and are specific to the parasite actin filaments.

## Results and discussion

### Plasmodium CPs form similar-shaped homo- and heterodimers in solution

*Pb*CPαα without the α-tentacle (*Pb*CPαα^ΔC20^) forms dimers with the canonical mushroom shape of CPs (24), but it was not clear whether the full-length *Pb*CPαα also could homodimerize in a similar manner. Here, we set out to investigate the quaternary arrangements of the full-length homo- and heterodimeric *Pb*CPs as well. Using a combination of small-angle X-ray scattering (SAXS) and homology modeling based on related crystal structures, we show that, indeed, all three parasite CP versions (the truncated and full-length *Pb*CPαα and *Pb*CPαβ) form similar pseudo-symmetric dimers in solution (**Fig. 1**). The SAXS data show folded, globular proteins with flexible parts, *Pb*CPαβ being somewhat more compact than the homodimers (**Fig. 1A and B**). Interparticle distances are similar for all PbCPs, with the heterodimer showing slight bimodality (**Fig. 1C**). Modeling into the SAXS data suggests that *Pb*CPαβ forms a tight structure, whereas the homodimers adopt a looser structure, which may arise from steric hindrances at the dimer interface (**Fig. 1D**). The orientation of a *Plasmodium-specific* insert of *Pb*CPα (24) and the tentacle domains of both subunits cannot be reliably deduced from the SAXS data, indicating that they are disordered in solution, which is characteristic of the tentacles in canonical CPs as well (13, 14).

**Fig. 1.**
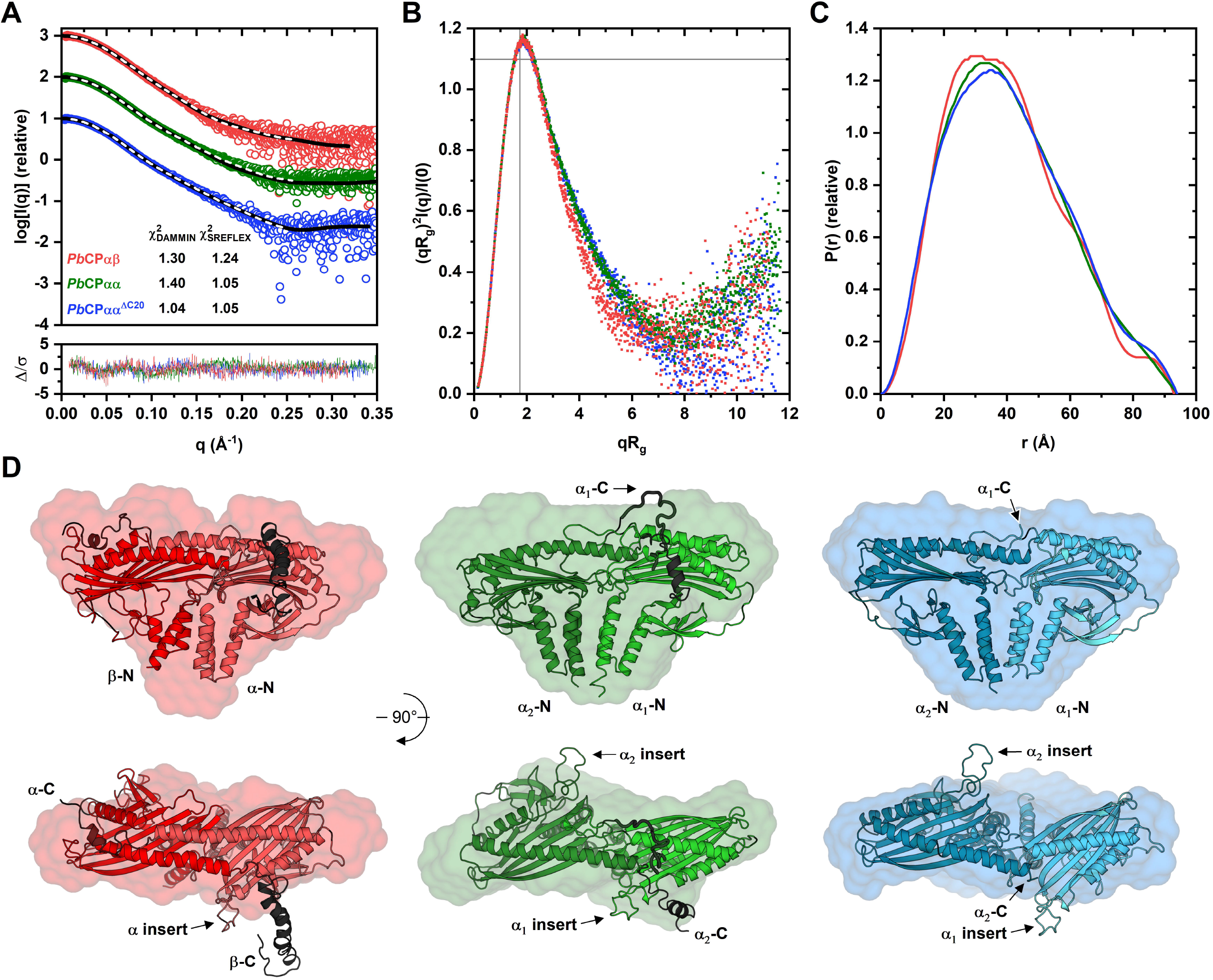
*Pb*CP homo- and heterodimers have a canonical shape in solution. (A) Experimental scattering curves of *Pb*CPαβ, *Pb*CPαα, and *Pb*CPαα^ΔC20^ (red, green, and blue open circles, respectively) and the respective fits of *DAMMIN* (white dashed lines) and *SREFLEX* (black line) models to the data. The χ^2^ values of the different models to the data are indicated. Weighted residuals of the *SREFLEX* model fits are denoted in the lower graph (lines colored respectively), where Δ/□ = [□_□□□_ (□) − □□_□□□_ (□)]□(□). (B) Dimensionless Kratky plot (colored similarly to panel A). (C) Real-space distance distribution plot (colored similarly to panel A). (D) Cartoon models of *Pb*CPs (colored as in panel A, with chain A in lighter color) and the corresponding SAXS envelopes as surface representations. The tentacle domains of each structure are colored black. Experimental data for *Pb*CPαα^ΔC20^ for comparison were used from a previous publication (24).

### Crystal structure of the *Pb*CPαα homodimer

To gain further structural insight into the unusual homodimerization of the parasite CP, we crystallized and determined the structure of the truncated *Pb*CPαα^ΔC20^ (**Fig. 2A**), as the full-length protein as well as the heterodimer have so far resisted crystallization. Extensive attempts to solve the structure using various molecular replacement approaches failed, pointing towards significant subunit- and domain-wise structural rearrangements in *Pb*CPαα^ΔC20^ compared to canonical CP heterodimers. We finally succeeded in solving the structure to 2.2 Å resolution using single-wavelength anomalous diffraction of bromide derivatives (**Table I**). As expected from SAXS analysis from before (24) and in this work, the two *Pb*CPα^ΔC20^ subunits form a less compact structure than the canonical heterodimer (**Fig. 2A**), still closely resembling the established mushroom shape of metazoan CPs (13, 14), despite the low sequence conservation (23). Albeit being a homodimer, the structure is not completely symmetric. As predicted by homology modeling and SAXS data, significant structural rearrangements are required at the dimer interface to accommodate two identical subunits in the dimer (**Fig. 2B**). This results in a new type of non-canonical dimeric CP structure and a new example of rare asymmetric homodimers among proteins in general (25). Because of the asymmetry, we will here refer to chain A of the crystal structure as *Pb*CPα_1_^ΔC20^ and chain B as *Pb*CPα_2_^ΔC20^. While in canonical heterodimers the stalk domains of the subunits pivot relative to the core structure to accommodate denser packing, in *Pb*CPαα^ΔC20^, the orientation is the same in both subunits, disrupting the compact structure (**Fig. 2C**). Further major structural deviations include longer helices of the stalk domains, a *Plasmodium-specific* insert between the globule and cap β-sheets (Ser135-Ala159), and a unique β-hairpin-like C-terminus (Leu268-Leu286) of the *Pb*CPα_1_^ΔC20^ H5 helix (**Fig. 2**).

**Fig. 2.**
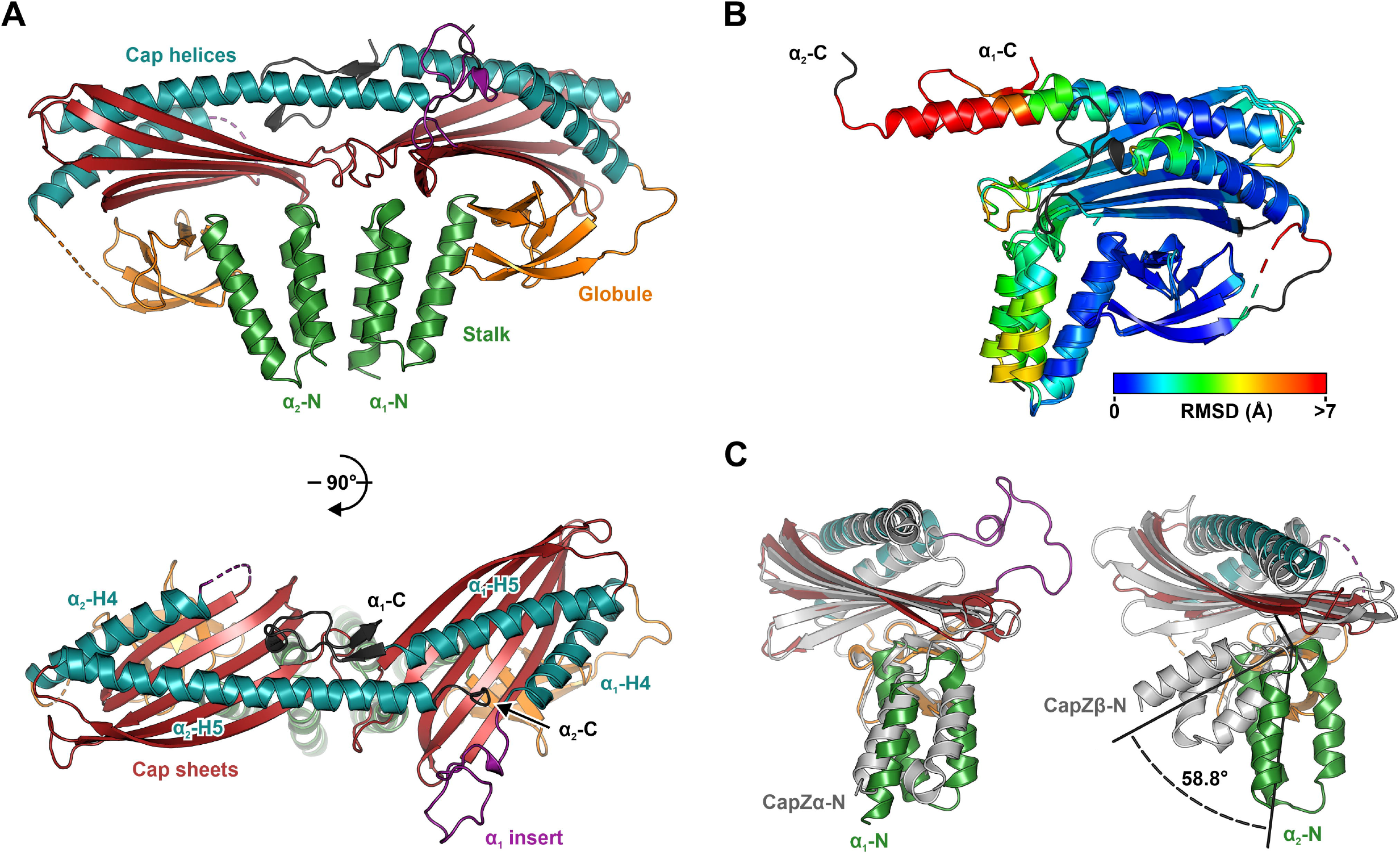
Crystal structure and domain arrangement of *Pb*CPαα^ΔC20^. (A) Crystal structure of *Pb*CPαα^ΔC20^. (B) Superposition of the *Pb*CPα_1_^ΔC20^ and *Pb*CPα_2_^ΔC20^ subunits. The rainbow color scale represents the r.m.s.d. between the subunits. Residues excluded from the alignment are colored black. (C) Superposition of *Pb*CPα_1_^ΔC20^ to CapZα (left) and *Pb*CPα_2_^ΔC20^ to CapZβ (right). The domains of *Pb*CPαα^ΔC20^ are colored as in panel A. CapZαβ [PDB ID: 1IZN (13)] is shown in light gray.

**Table I.**
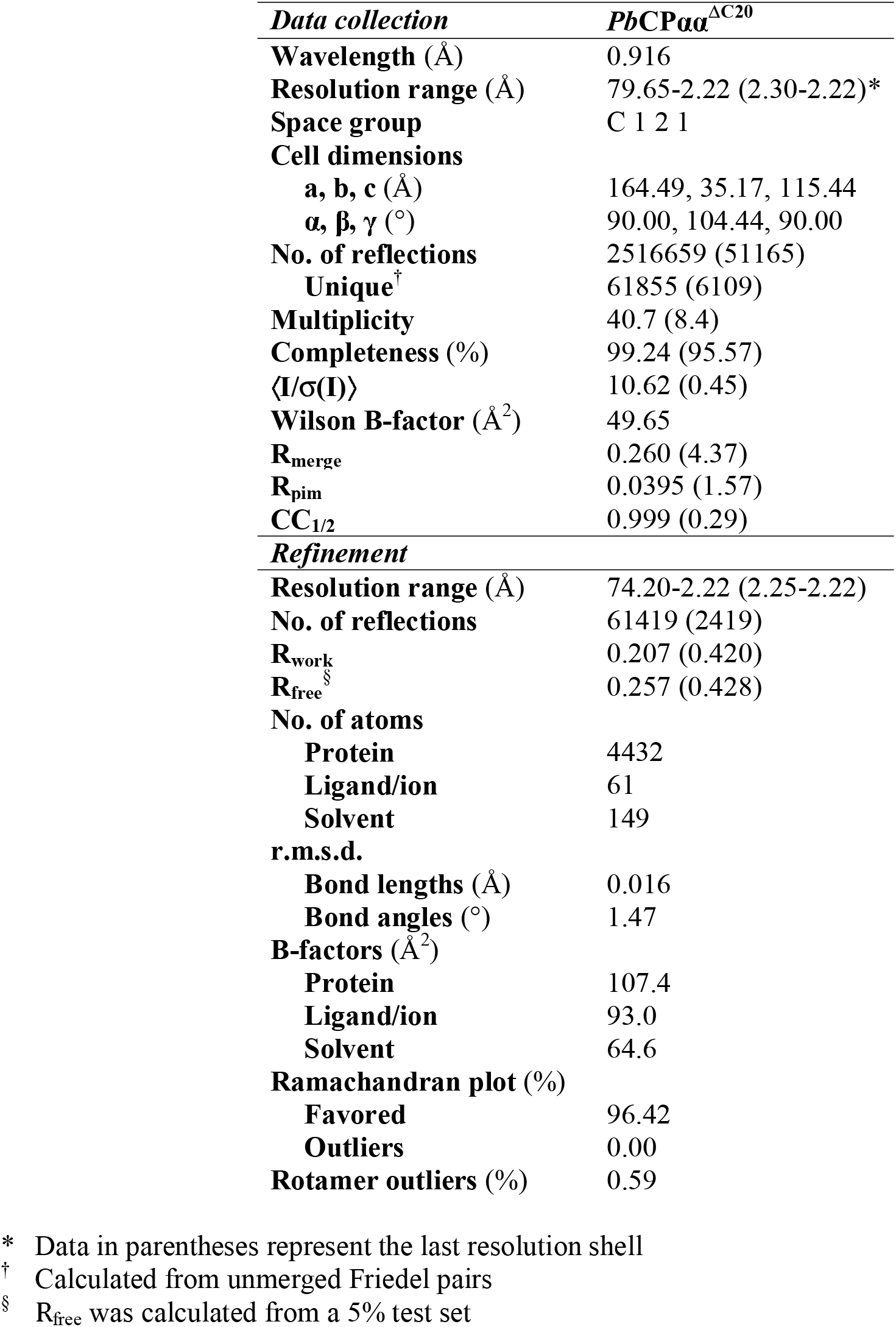
Data collection and refinement statistics.

| *Data collection* | *PbCPaa*<sup>ΔC20</sup> |
| --- | --- |
| **Wavelength** (Å) | 0.916 |
| **Resolution range** (Å) | 79.65-2.22 (2.30-2.22)* |
| **Space group** | C 1 2 1 |
| **Cell dimensions** |  |
| **a, b, c** (Å) | 164.49, 35.17, 115.44 |
| **α, β, γ** (°) | 90.00, 104.44, 90.00 |
| **No. of reflections** | 2516659 (51165) |
| **Unique**<sup>†</sup> | 61855 (6109) |
| **Multiplicity** | 40.7 (8.4) |
| **Completeness** (%) | 99.24 (95.57) |
| **⟨I/σ(I)⟩** | 10.62 (0.45) |
| **Wilson B-factor** (Å<sup>2</sup>) | 49.65 |
| **R<sub>merge</sub>** | 0.260 (4.37) |
| **R<sub>pim</sub>** | 0.0395 (1.57) |
| **CC<sub>1/2</sub>** | 0.999 (0.29) |
| **Refinement** |  |
| **Resolution range** (Å) | 74.20-2.22 (2.25-2.22) |
| **No. of reflections** | 61419 (2419) |
| **R<sub>work</sub>** | 0.207 (0.420) |
| **R<sub>free</sub>**<sup>§</sup> | 0.257 (0.428) |
| **No. of atoms** |  |
| **Protein** | 4432 |
| **Ligand/ion** | 61 |
| **Solvent** | 149 |
| **r.m.s.d.** |  |
| **Bond lengths** (Å) | 0.016 |
| **Bond angles** (°) | 1.47 |
| **B-factors** (Å<sup>2</sup>) |  |
| **Protein** | 107.4 |
| **Ligand/ion** | 93.0 |
| **Solvent** | 64.6 |
| **Ramachandran plot** (%) |  |
| **Favored** | 96.42 |
| **Outliers** | 0.00 |
| **Rotamer outliers** (%) | 0.59 |

\* Data in parentheses represent the last resolution shell
<sup>†</sup> Calculated from unmerged Friedel pairs
<sup>§</sup> R<sub>free</sub> was calculated from a 5% test set

Due to the reduced number of intersubunit and crystal contacts, *Pb*CPα_2_^ΔC20^ is more disordered, with a higher average B-factor than *Pb*CPα_1_^ΔC20^ (**Suppl. Fig. S1 and Table I**), similar to what has been described for CapZαβ (14). Consequently, the *Plasmodium-specific* insert (residues 134-155) is also not visible in *Pb*CPα_2_^ΔC20^. In *Pb*CPα_1_^ΔC20^, this loop protrudes from the structure, making space for the long helix H5 (residues 243-283) of *Pb*CPα_2_^ΔC20^, which would precede the α-tentacle in the full-length protein. In *Pb*CPα_1_^ΔC20^, H5 is broken at residue 267 and turns back towards the same subunit in a β-hairpin-like structure, instead of extending into *Pb*CPα_1_^ΔC20^. The base of the *Plasmodium-specific* loop seems to be stabilized by a partial disulfide bond between Cys158 and Cys179 in the *Pb*CPα_1_^ΔC20^ subunit but not in *Pb*CPα_2_^ΔC20^. The loose nature of the homodimer and the substantial structural differences in the C-terminal region could result in a non-canonical binding mode to the highly dynamic *Plasmodium* actins (26, 27) and a distinct role for the homodimers as compared to the canonical or the parasite CP heterodimers (19, 23, 24).

In comparison to the sarcomeric [PDB ID: 1IZN (13)], cytoplasmic [PDB ID: 4AKR (14)], and dynactin bound [PDB ID: 6F1T (28)] CP isoforms, *Pb*CPαα^ΔC20^ has an increased surface area but significantly decreased dimer interface area (**Suppl. Table SI**), as is typical for asymmetric homodimers (25). Root-mean-square deviations (r.m.s.d.) calculated between these structures reveal further insights on a dimer, subunit, and domain level (**Suppl. Table SII**). Canonical heterodimers are much more similar to each other than to the *Plasmodium* homodimer (**Suppl. Table SIIA**), owing to their subunit arrangement. CP-related CATH domains (1.20.1290.20 and 2.40.160.80) are present in *Pb*CPαα^ΔC20^, manifested by a lower r.m.s.d. when individual subunits are compared. Both *Pb*CPαα^ΔC20^ subunits are more similar to canonical CPα, with *Pb*CPα_2_^ΔC20^ being slightly closer to CPβ (**Suppl. Table SIIB**). Given sequence limitations, this subunit could facilitate a more canonical heterodimer arrangement of the *Plasmodium* homodimer. Individual domains of *Pb*CPαα^ΔC20^ fit well to canonical CP subunits suggesting the importance of individual structural elements in the capping function (**Suppl. Table SIIC**), despite sequence differences.

### PbCPs control actin polymerization in a non-canonic manner

CPs typically block the barbed end, increasing its apparent critical concentration (Cc_app_) to that of the pointed end’s (2). *Pb*CPs increase the supernatant fraction of *Pf*ActI in pelleting assays (24), which could either be explained by filament shortening or by limited depolymerization due to the increase of Cc_app_. In dilution series of fluorescently labeled *Pf*ActI filaments, none of the *Pb*CPs affect Cc_app_, even at high concentrations (**Fig. 3A**). Furthermore, gelsolin, a major barbed end capper (1) absent from *Plasmodium* spp. (18), does not affect the Cc_app_ of *Pf*ActI filaments (**Fig. 3B**) or *Pf*ActI depolymerization dynamics (27). None of the *Pb*CPs increase the Cc_app_ of heterologous skeletal α-actin either (**Fig. 3C**), despite them reportedly being able to modulate α-actin (22) or non-muscle β-actin polymerization (19, 23). We hypothesize that *Pb*CPs have a lower affinity to actin than canonical CPs, allowing actin subunit exchange, making them wobbly or leaky cappers (2), similar to formins (1).

**Fig. 3.**
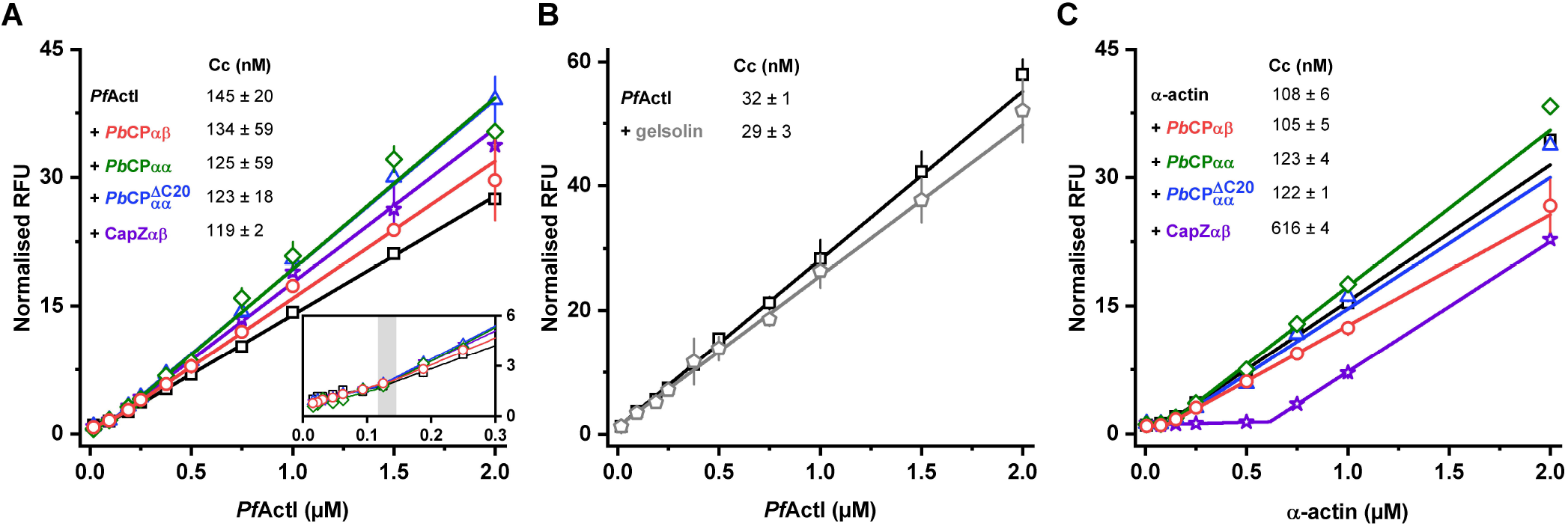
*Pb*CPs do not affect the critical concentration of actin polymerization. (A) *Pf*ActI filaments (open black squares) capped by *Pb*CPαβ, *Pb*CPαα, *Pb*CPαα^ΔC20^, and CapZαβ (open red circles, green diamonds, blue triangles, and purple stars, respectively). The gray shading indicates the range of determined Cc_app_ from two-line fits (respectively colored lines). (B) Cc plot of *Pf*ActI filaments (open black squares) with gelsolin (open gray pentagon) under Ca^2+^ conditions. (C) α-actin filaments (open black squares) capped by *Pb*CPαβ, *Pb*CPαα, *Pb*CPαα^ΔC20^, and CapZαβ (as in panel A). Errors represent SD (n = 3). RFU = relative fluorescent unit, normalized to the lowest concentration.

The inability of *Pb*CPs to increase Cc_app_ raises a question about the presence of other actin regulatory properties. In higher eukaryotes, CPs nucleate filaments, abolishing the lag phase, but block subunit exchange at the barbed end (2), reducing the initial elongation velocity and steadystate filament mass. In a homologous system, contrary to expected, we found that *Pb*CPs increase elongation velocity and considerably raise the steady-state level of F-actin mass (**Fig. 4A**). Their effect on nucleation is ambiguous due to the nature of *Pf*ActI polymerization curves (27). In preseeded assays, the filament mass is less affected (**Fig. 4B**). CapZαβ does not seem to modulate *Pf*ActI polymerization notably. On the contrary, both *Pb*CP homodimers and the heterodimer behave similarly to typical CPs with α-actin (**Fig. 4C and D**). These results suggest that *Pb*CP homo- and heterodimers have similar, redundant functions, which would secure actin capping functions in the blood stages of the parasite where *Pb*CPβ is phenotypically absent (19). *Pb*CPαα^ΔC20^ displays a diminished, yet comparable capping effect, despite the absence of the essential tentacle domains (15, 16). The reduced importance of the tentacle domain in *Pb*CPs has been suggested before (24). However, *in vivo*, the *Plasmodium* α-tentacle seems to be indispensable (23). The major difference between homo- and heterodimers is seen in the initial filament mass in non-seeded systems (**Fig. 4A and C**). It seems that while *Pb*CPs are able to cap F-actin, in the absence of preformed filaments, the homodimers may also be able to stabilize or sequester short, non-nucleating structures, perhaps lateral dimers (27). This behavior of the homodimers extends to α-actin and seems to be independent of the presence of the α-tentacle.

**Fig. 4.**
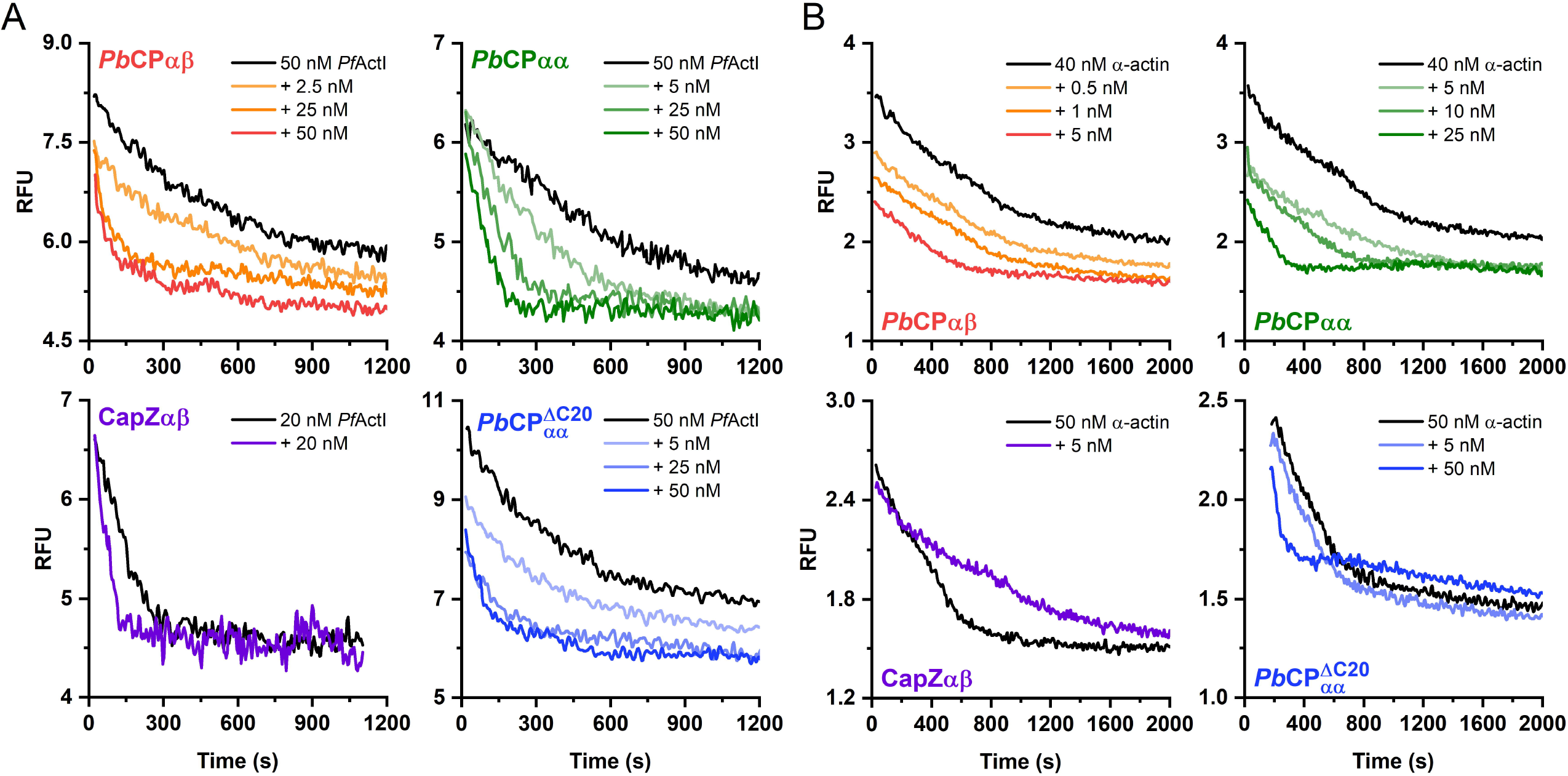
Atypical regulation of polymerization kinetics by *Pb*CPs. (A) Polymerization curves of *Pf*ActI in the absence and presence of increasing concentrations of *Pb*CPαβ, *Pb*CPαα, *Pb*CPαα^ΔC20^, and CapZαβ. (B) Polymerization of *Pf*ActI on capped, preformed homologous filaments in the absence and presence of increasing concentrations of *Pb*CPαβ, *Pb*CPαα, *Pb*CPαα^ΔC20^, and CapZαβ. (C and D) Non-seeded and seeded polymerization curves of α-actin with the different CPs, as in panels A and B. RFU = relative fluorescent unit.

### Dilution-induced disassembly of filaments is enhanced in the presence of PbCPs

*Pf*ActI filaments are highly dynamic (26, 27) with high disassociation rates at both ends (29). Upon dilution below their Cc_app_, filaments decompose rapidly, faster than α-actin (27). We measured the barbed end blocking efficiency of *Pb*CPs, by following the disassembly of fluorescently labeled *Pf*ActI and α-actin filaments. *Pb*CPs facilitate depolymerization of both *Pf*ActI (**Fig. 5A**) and α-actin filaments (**Fig. 5B**). While CapZαβ behaved as expected with α-actin, reducing the rate of depolymerization (2), it also increased the rate of *Pf*ActI depolymerization, suggesting that the effect results from an interplay between *Plasmodium* actins and CPs, not strictly from the latter. While the suggested propensity of *Pb*CPs to sequester actin dimers agrees well with the data, we cannot exclude a moonlighting function of these proteins as filament severing proteins (1). Such behavior of CPs has been described in the presence of VASP proteins (30). *Pb*CPs, however, in the absence of ENA/VASP homologs (18), might have evolved to have inherent severing capability. This would raise the question of redundancy among the limited *Plasmodium* actin regulators, as multiple severing proteins have been described (17, 27).

**Fig. 5.**
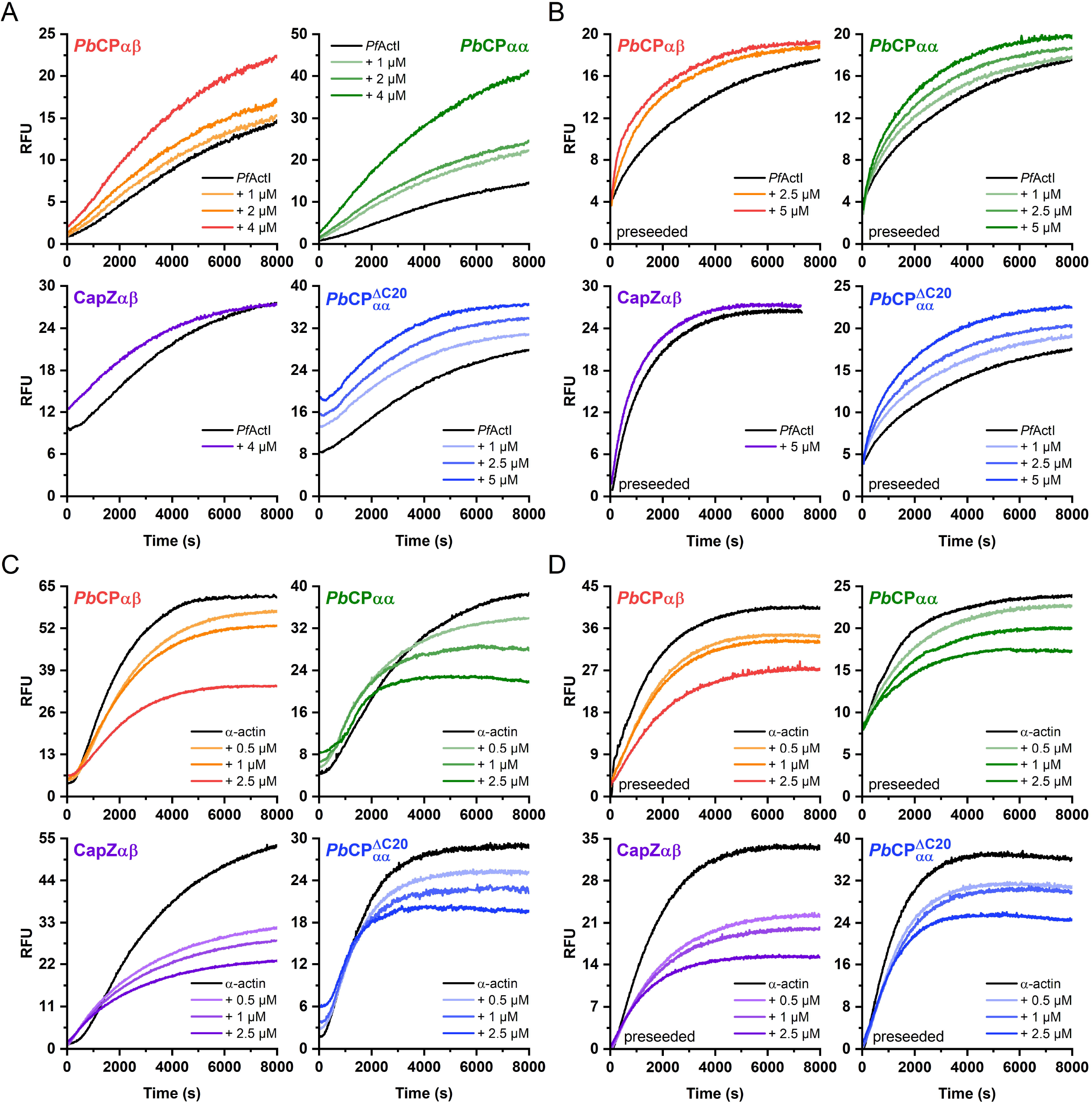
*Pb*CPs facilitate filament depolymerization. (A) Dilution-induced depolymerization of *Pf*ActI filaments (black line) with various concentrations of *Pb*CPαβ, *Pb*CPαα, *Pb*CPαα^ΔC20^, and CapZαβ. (B) Depolymerization of *Pb*CPαβ, *Pb*CPαα, *Pb*CPαα^ΔC20^, and CapZαβ capped α-actin filaments (black line). RFU = relative fluorescent unit.

### Implications of an atypical CP

*In vivo*, many actin binding proteins (ABPs) exist to fine-tune barbed end capping through well-characterized mechanisms (2). While the majority of these are absent from *Plasmodium* (17, 18), we cannot exclude the possibility of other, so far uncharacterized, regulators. Our search for V-1/myotrophin homologs and the CP-binding and uncapping motif among known *Plasmodium* transcripts or for the S100B interaction motif in *Plasmodium* CPs resulted in no clear hits. To date, in *P. knowlesi*, PIP2 is the only described direct CP regulator, with HSC70 mentioned as a binding partner, uninvolved in capping activity (22). Residues involved in canonical interaction with proteinaceous regulators are generally not conserved in *Plasmodium* (**Suppl. Fig. S2 and S3**), with an exception of several basic residues (K278 and K282 in *Pb*CPα, R233 in *Pb*CPβ) involved in canonical F-actin interaction and PIP2 binding (15, 16, 31). Based on our kinetic data, *Pb*CPs seem to have at least an order of magnitude lower affinity towards *Pf*ActI filaments compared to CapZαβ and canonical actin. Left unregulated, CPs would constantly cap actin filaments, completely arresting cell motility (10). We cannot rule out that this innate down-regulation of *Pb*CPs could allow for the existence of free barbed ends in the absence of other regulators, allowing the parasite to function with only a minimal set of ABPs.

Given the discussed substantial differences of *Pb*CPs compared to canonical CPs, in particular the absence of a β-subunit in certain stages, we hypothesize that the F-actin binding mode of homodimeric *Pb*CPs may also be different. Canonical CPs bind the barbed end through an electrostatic interaction mediated largely by the basic triad on CapZαβ and an acidic patch on the last two actin protomers (15). To understand whether this binding mode can exist in *Plasmodium*, we used molecular docking to generate a model of a *Pb*CPαα^ΔC20^ capped *Pf*ActI filament (**Fig. 6A**). Presence of Arp1 in the *P. berghei* genome (32) could implicate the existence of *Pb*CP capped Arp1 filaments in *Plasmodium*, thus allowing the cryo-EM structure of CapZαβ capped Arp1 filament in *Sus scrofa* dynactin (28) to be used as basis for further modeling studies. Electrostatic potential surfaces of the barbed end (**Fig. 6B**) and CPs (**Fig. 6C**) reveal the absence of canonical charged interaction points only for *Plasmodium* (**Suppl. Fig. S4)**, even though residues involved in forming these interaction surfaces are conserved (**Suppl. Fig. S2, S3, and S5**). In *Pf*ActI, flanking residues mask the apparent charge of the patch (**Fig. 6B**). Differences in electrostatic potential surfaces (33) could also play a large role in the divergent nucleation and polymerization properties of *Pf*ActI (27). Residues of the basic triad in *Pb*CPαα^ΔC20^ are more buried in the structure, not in close proximity to each other and the slight asymmetry of the homodimer does not compensate significantly for the loss of contributing residues of *Pb*CPβ. Possibly due to the absence of CARMIL proteins in *Plasmodium*, a positively charged canonical binding cavity is not present in *Pb*CPαα^ΔC20^ (**Fig. 6C**). In the CapZαβ heterodimer, the β-tentacle locks the complex by binding the hydrophobic pocket of the terminal actin subunit (15, 16). In our model, the *Plasmodium-specific* insert of *Pb*CPα_1_^ΔC20^ is close to the expected position of the β-tentacle. This insert shares many possible, hydrophobic or electrostatic, interaction points with the β-tentacle (**Fig. 6D**), residues mainly conserved in *Plasmodium* (**Suppl. Fig. S2**), suggesting a role for the insert in barbed end binding in the absence of canonical interaction motifs. Mapping sequence conservation of *Plasmodium* CPα sequences on the structure of *Pb*CPαα^ΔC20^ reveals that core residues, especially the ones involved in the intersubunit surface are highly conserved, suggesting that other CPαα homodimers could exist in *Plasmodium* spp. Canonical binding partner interaction points present in CapZαβ lack localized sequence conservation in *Pb*CPαα^ΔC20^ (**Fig. 6E and Suppl. Fig. S2**).

**Fig. 6.**
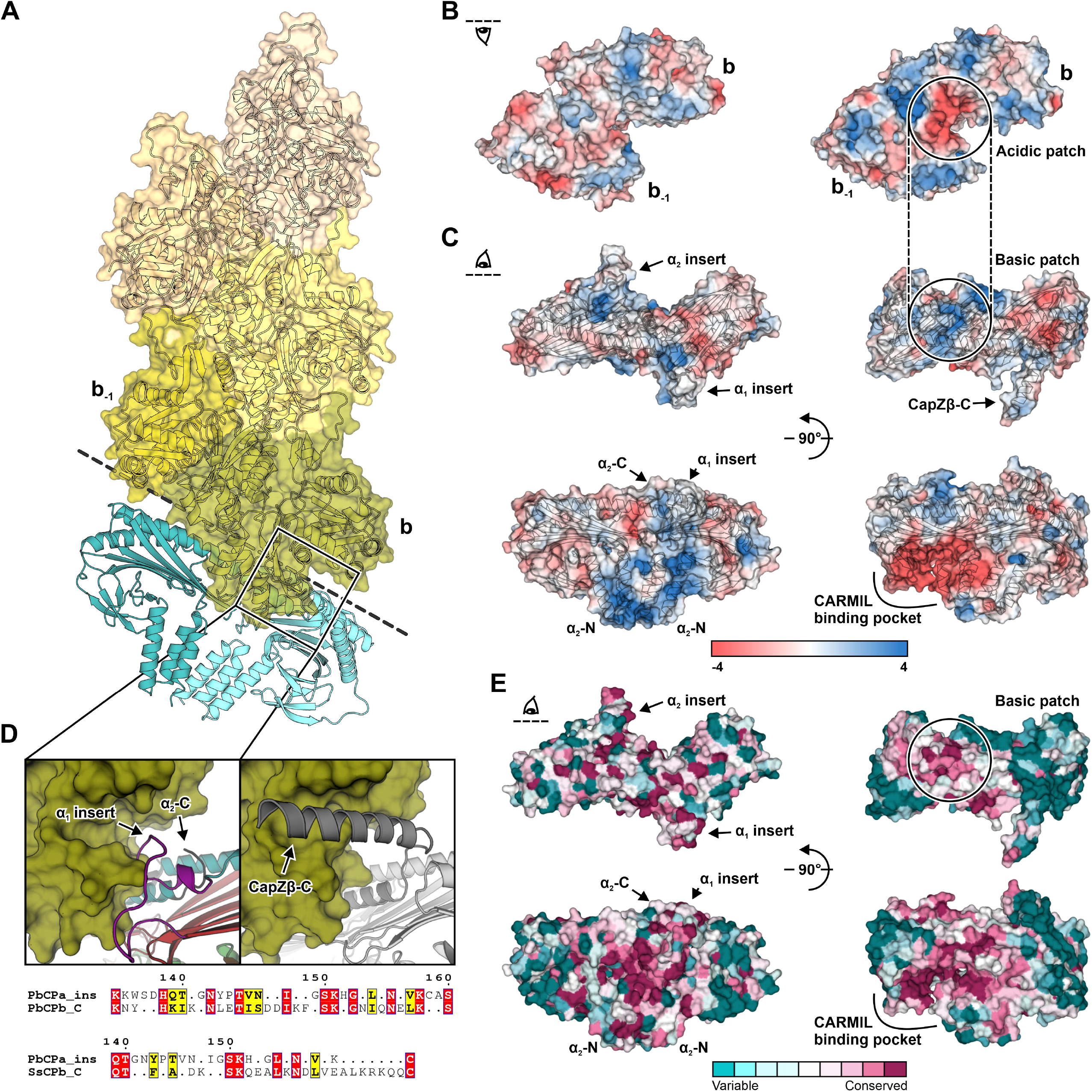
Atypical structural properties of *Pf*ActI and *Pb*CPαα^ΔC20^ suggest new barbed end binding modes. (A) Model of *Pf*ActI filament [green to beige colored surfaces, PDB ID: 5OGW (51)] capped by *Pb*CPαα^ΔC20^ (blue cartoon) in a canonical arrangement. (B) Electrostatic potential surface of the last (b) and penultimate (b-1) protomer of *Pf*ActI (left) and Arp1 [right, PDB ID: 6F1T (28)] filament barbed ends. (C) Electrostatic potential surface of *Pb*CPαα^ΔC20^ (left) and CapZαβ [right, PDB ID: 6F1T (28)]. (D) The hydrophobic pocket of the last subunit of *Pf*ActI (left) and Arp1 (right) filament, represented as green surface. The *Plasmodium-specific* insert, cap sheets, and cap helices of *Pb*CPαα^ΔC20^ (left) are colored purple, red, and teal, respectively. The CapZαβ (right) α- and β-subunits are shown in light and dark gray, respectively. Sequence alignment (bottom) of the *Plasmodium* insert (PbCPa_ins) with the β-tentacles of *P. berghei* (PbCPb_C: A0A509AQN8) and *S. scrofa* CP (SsCPb_C: A9XFX6), grouped and colored respective to a Risler matrix, using the *ESPript* convention. (E) Structure of *Pb*CPαα^ΔC20^ (left) and CapZαβ (right) with the surface colored by *ConSurf* residue conservation scores from low (cyan) to high (magenta).

The truncated *Pb*CPαα^ΔC20^ binds actin in the absence of the essential structural elements (24), suggesting that canonical functions of the tentacle domains are compensated by other means. Possibilities are that both interaction sites undergo significant conformational changes upon binding, or a completely novel mechanism is involved. Follow-up mutagenesis studies aimed at the *Plasmodium* insert and the surface residues of the capping structure could shed light to this unique interaction.

## Concluding remarks

CPs are essential regulators of the cytoskeleton and conserved among metazoans as heterodimeric proteins, which are important *e.g*. for lamellipodial protrusions at the leading edge of crawling cells (4). Apicomplexan parasites display fast gliding motility in the absence of changes in cell shape (34). It seems that due to different needs during different lifecycle stages, these parasites have evolved an additional unique homodimeric form of CP (24) as part of their limited repertoire of ABPs (17). In contrast to canonical CPs (2), *Pb*CPs facilitate the rapid turnover of the dynamic *Plasmodium* actin filaments. Due to the atypical structural and biochemical properties, and a unique mode of binding to the barbed ends as well as their essential nature, *Plasmodium* CPs could prove as promising drug targets against malaria.

## Materials and methods

### Protein expression and purification

*Pb*CPαβ, *Pb*CPαα, *Pb*CPαα^ΔC20^, CapZαβ, α-actin, and *Pf*ActI were prepared as described previously (24, 27). Human full-length gelsolin was purchased from Cytoskeleton (US).

### Model refinement using small-angle X-ray scattering (SAXS)

SEC-SAXS data of *Pb*CPαα and *Pb*CPαβ, at a respective concentration of 12 and 5.8 mg/ml, were collected on the B21 beamline at Diamond Light Source (Didcot, UK). Data processing, *ab initio DAMMIN* (35) modeling, and visualization were carried out as previously described (24). Initial models of the homodimers were prepared by extending the crystal structure of *Pb*CPαα^ΔC20^ using *SWISS-MODEL* (36). The tentacle domain of *Pb*CPα_1_ was modeled using *EOM*(37). A model of *Pb*CPαβ was assembled from the modeled *Pb*CPα_2_ subunit and *I-TASSER* (38) modelled *Pb*CPβ monomer using the structure of dynactin-bound *S. scrofa* CapZαβ [PDB ID: 6F1T (28)] as a template. The models were corrected against major steric clashes using *Coot* (39) and energy minimized using *UCSF Chimera* (40). The *Pb*CP models were split into domains and refined with normal mode analysis using *SREFLEX* (41).

### Crystallization and structure solution

*Pb*CPαα^ΔC20^ was crystallized at 4°C using the vapor diffusion method. Crystals were grown from a 1:1 drop ratio of 10-15 mg/ml protein and precipitant [100 mM MES pH 6.5, 100-150 mM triammonium citrate, 10-12% (w/v) PEG 20000, and 0.5 M NDSB-195 (Hampton, US)] and subsequently improved by iterative micro-seeding. The crystals were soaked for 1-2 min in 1 M NaBr before flash freezing in 15% (v/v) glycerol. Diffraction datasets were collected on the I04-1 beamline at Diamond Light Source (Didcot, UK) using a Pilatus 6M detector and 0.916 Å beam wavelength. Datasets from multiple crystals were integrated, scaled, and merged using *DIALS* (42) in the *xia2* pipeline (43). Experimental phases and an initial main-chain trace were obtained from *SHELX* (44). The structure was refined using the *CCP4* package (45) and *Phenix* (46), with iterative rebuilding in *Coot*.

### Structural bioinformatics

Details of the surface area and r.m.s.d. calculations are found in **Supplementary Information**. Database searches against homologs and motifs were carried out using the PlasmoDB database (47), *ScanProsite* (48), *MyHits* (49), and *PATTINPROT* (50). Model of *Pb*CPαα^ΔC20^ capped *Pf*ActI filament was prepared by aligning 5 protomers of *Pf*ActI filament [PDB ID: 5OGW (51)] and *Pb*CPαα^ΔC20^ to the CapZαβ capped barbed end of the Arp1 filament in dynactin [PDB ID: 6F1T (28)] using *TM-align* (52). Gaps in the structure of *Pb*CPαα^ΔC20^ were modeled in using *SWISS-MODEL*. Relative orientation of *Pb*CPαα^ΔC20^ was refined with *RosettaDock* (53), while residues involved in the interface were energy minimized with *UCSF Chimera*. Electrostatic potential surfaces were calculated using *APBS* (54). *EMBOSS Matcher* (55) was used for local sequence alignments and *ESPript* (56) for their visualization. 21 non-redundant *Plasmodium* CPα sequences from PlasmoDB were analyzed, with conservation score subsequently mapped on the structure of *Pb*CPαα^ΔC20^ using *ConSurf* (57). CapZαβ was prepared similarly, using automatically retrieved homolog sequences.

### Actin-CP interaction assays

Fluorescence-based polymerization assays were carried out in triplicate and analyzed as previously described (27) with minor modifications. Cc_app_ was determined using a dilution series prepared from 10 μM actin polymerized together with either 10 μM CP or 0.4 μM gelsolin for *Pf*ActI, and 0.4 μM CP for α-actin, respectively. EGTA was omitted from the buffers for the samples with gelsolin. In the polymerization assays, the total actin concentration was kept at 4 μM. 2 μM α-actin, or 0.5 μM *Pf*ActI filaments (2 μM in the CapZαβ assay) were used as nuclei in preseeded polymerization assays. In the depolymerization assays, CPs were incubated with 2-5 μM actin filaments for 5 min at 20°C prior to 100-fold dilution. The polymerization data were despiked before averaging. No pointed end cappers were used in the assays to limit subunit exchange to the barbed end.

## Acknowledgements

We thank Dr. Herwig Schüler for providing the parasite CP cDNA constructs and Dr. Shuichi Takeda for the CapZαβ plasmid. The use of the facilities and expertise of the Biocenter Oulu Structural Biology core facility is gratefully acknowledged. We also thank the excellent beamline support at Diamond Light Source (B21 and I04-1) and the crystallographic community for their helpful comments on the structure of *Pb*CPαα^ΔC20^. This work was supported by the Academy of Finland, the Sigrid Juselius Foundation, and the Norwegian Research Council. The atomic coordinates and structure factors of *Pb*CPαα^ΔC20^ have been deposited in the Protein Data Bank, https://www.ebi.ac.uk/pdbe/ (PDB ID: 7A0H).

## Author contributions

I.K. and P.K. conceived the study. Á.Á.B. performed the experiments, analyzed the data, and drafted the manuscript. Á.Á.B., I.K. and P.K. solved and refined the crystal structure. Á.Á.B. and I.K. wrote the final manuscript and all authors approved the final version of the manuscript.

