## Supplemental data file for "Structure of an atypical homodimeric actin capping protein from the malaria parasite"

### SUPPLEMENTARY METHODS

#### Sequence alignment

Protein sequence alignments utilizing structural information were carried out using *T-COFFEE* (1). The alignments were visualized using *ESPript* (2). Secondary structure and residue contact information were extracted from the *SWISS-MODEL* (3) extended *Pb*CPαα$^{\Delta C20}$ structure and the *I-TASSER* (4) homology model of *Pb*CPβ for the CPα and CPβ subunits, respectively. *ConSurf* (5) conservation was calculated as discussed in **Materials and Methods**. Residues of canonical CPs involved in characterized interactions are reviewed in Eckert *et al*. (6). Residues involved in CP binding through forming the acidic or hydrophobic patch in actin or Arp1 are taken from literature sources (7, 8).

#### Electrostatic potential calculation

Surface potential of cytoplasmic β-actin and CapZαβ were calculated using *APBS* (9). A barbed end model was assembled by substituting the capped Arp1 filament in 6F1T (10) with β-actin present in the same entry and the CapZαβ with chains A and B from 1IZN (11). The protein chains were aligned using *TM-align* (12).

#### Surface area calculation

Gaps in the *Pb*CPαα$^{\Delta C20}$ structure were extended using *SWISS-MODEL* prior to analysis. The dimer interface and surface areas of the CPs were calculated using the *jsPISA* server (13). The total accessible surface area (ASA) for *Pb*CPαα$^{\Delta C20}$ and CapZαβ (PDB ID: 6F1T) were calculated using *PyMOL* (14).

#### Root-mean-square deviation calculation

Sequence-independent root-mean-square (r.m.s.d.) calculations of the CP structures and domains [PDB ID: 1IZN, 4AKR (6), 6F1T, and 7A0H (this work)] were carried out using *TM-align*. Gaps in the *Pb*CPαα$^{\Delta C20}$ structure were extended using *SWISS-MODEL* prior to analysis.

### SUPPLEMENTARY FIGURES AND TABLES

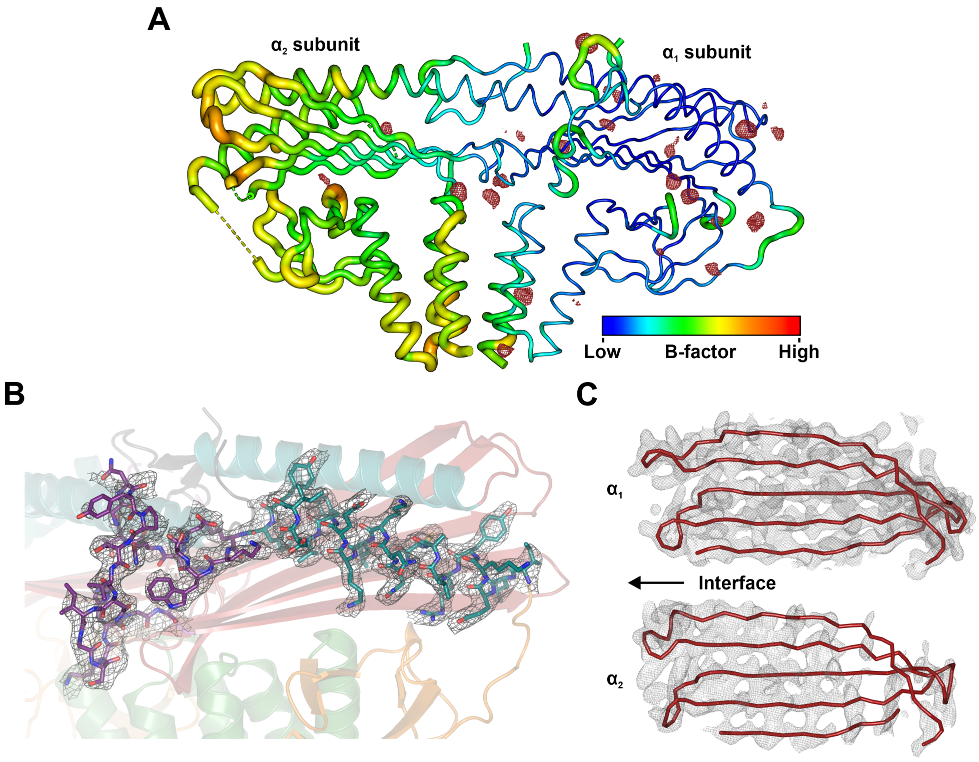


**Figure S1.** Loose dimer arrangement in *Pb*CPαα$^{\Delta C20}$. (A) *Pb*CPαα$^{\Delta C20}$ dimer colored according to B-factors with anomalous difference Fourier peaks of bromide atoms, contoured at 3 σ. Aspherical peak shapes suggest minor differences in bromide atom positions among merged datasets. (B) Electron density of the H4 helix and *Plasmodium* insert of *Pb*CPα$_1^{\Delta C20}$. (C) Electron density of the cap sheets of *Pb*CPα$_1^{\Delta C20}$ (top) and *Pb*CPα$_2^{\Delta C20}$ (bottom). The 2F$_o$-F$_c$ composite maps are contoured at 1 σ.

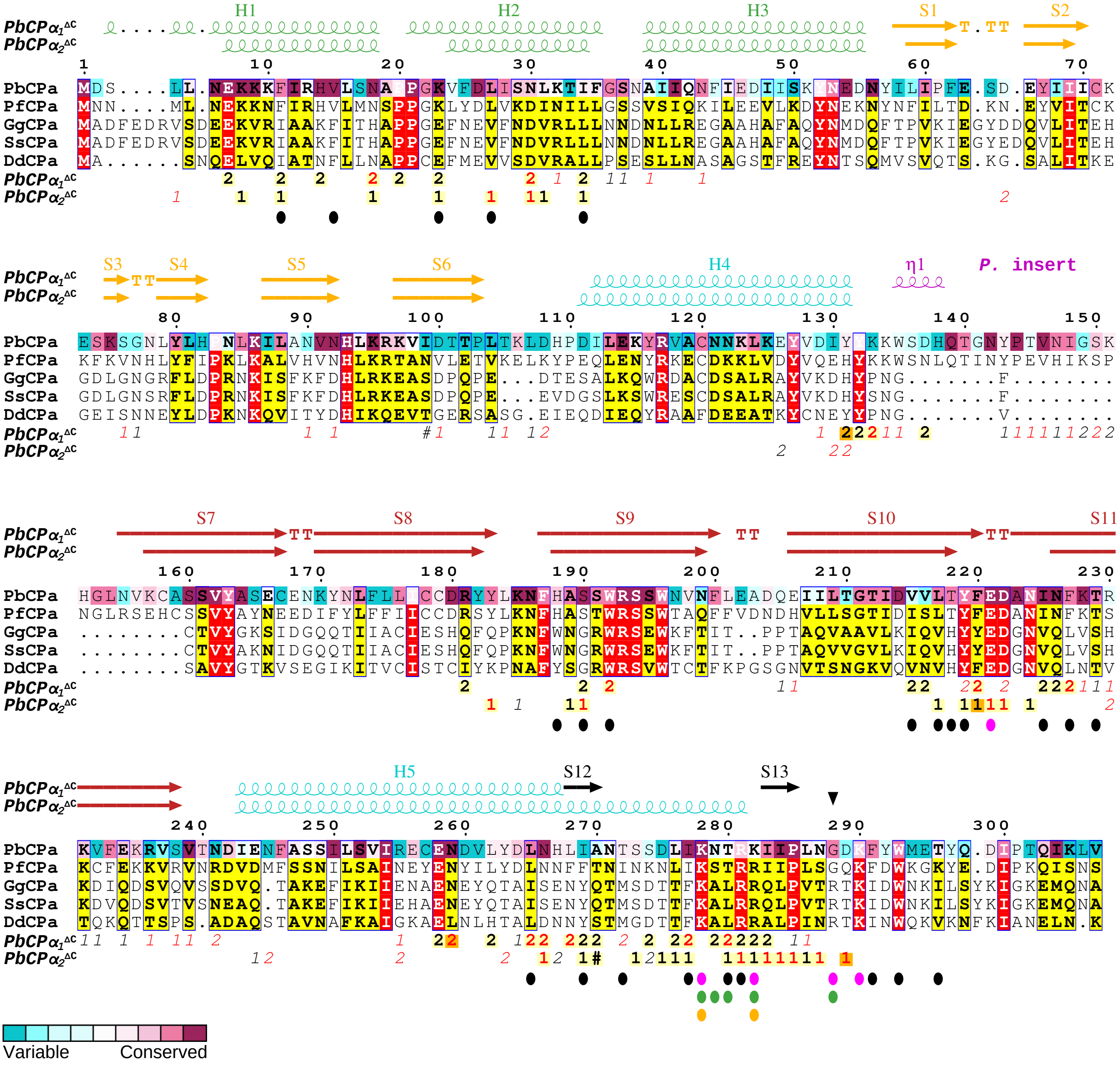


**Figure S2.** Sequence conservation of CPα. The secondary structure elements of *Pb*CPα$_1^{\Delta C20}$ and *Pb*CPα$_2^{\Delta C20}$ are indicated on top of the alignment and colored as follows: green for the stalk, orange for the globule, cyan for the cap helices, purple for the *Plasmodium* insert, red for the cap sheets, black for the C-terminus. The black triangle denotes the truncation point and start of the His$_6$-tag in *Pb*CPαα$^{\Delta C20}$ (15). The aligned sequences (PbCPa: A0A509AR49, PfCPa: Q8I3I2, GgCPa: P13127, SsCPa: A0PFK5, DdCPa: P13022) were grouped and colored respective to a Risler matrix, using the *ESPript* convention. The PbCPa sequence is colored by *ConSurf* conservation scores (low: cyan, high: magenta) among *Plasmodium* CPα sequences. Intersubunit and crystallographic contacts are shown below the sequences using the *ESPript* convention (1: *Pb*CPα$_1^{\Delta C20}$, 2: *Pb*CPα$_2^{\Delta C20}$). Residues important for canonical intraheterodimer contacts, actin-, V1/myotrophin-, and PIP$_2$-binding are indicated by black, magenta, green, and orange dots, respectively.

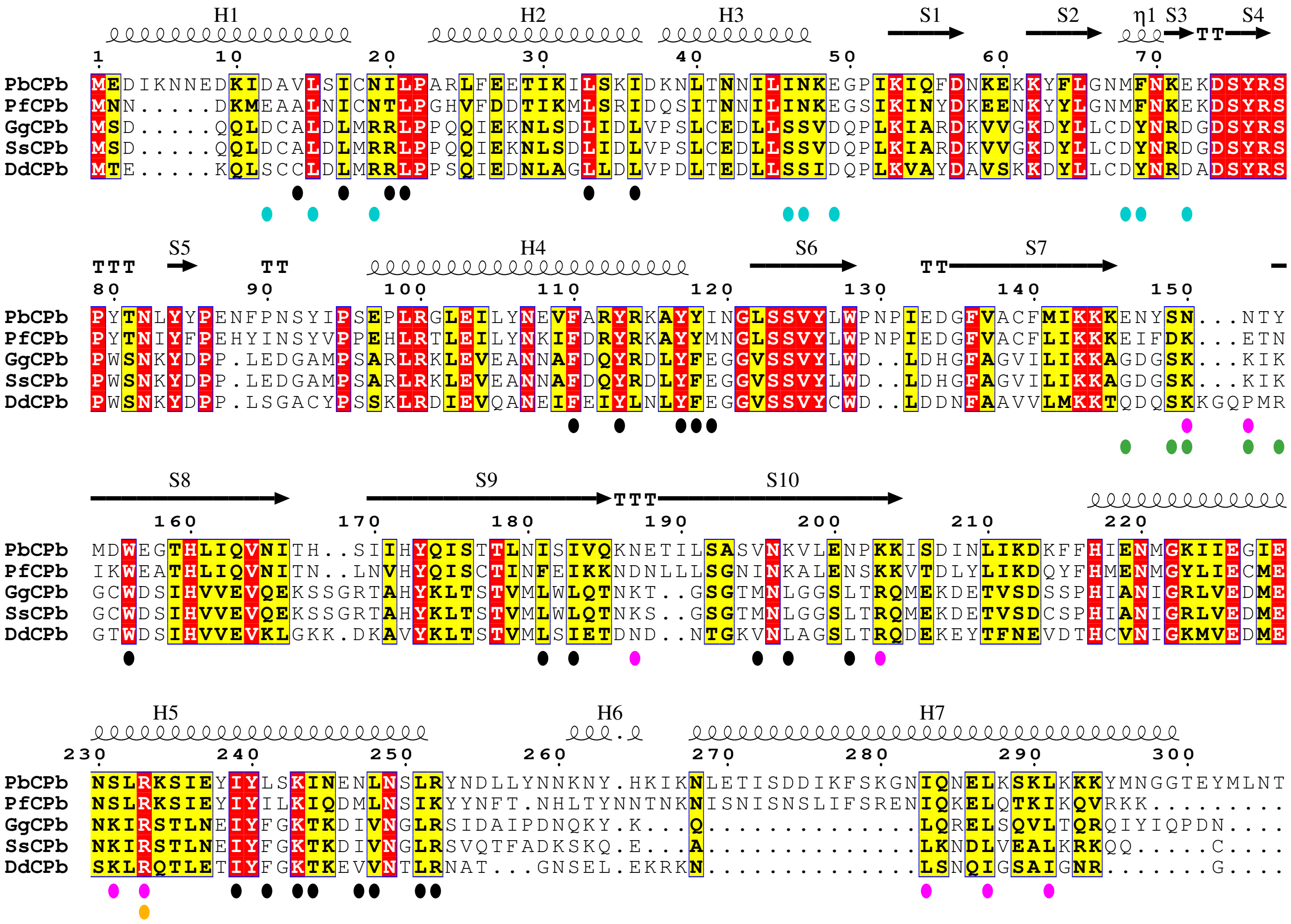


**Figure S3.** Sequence conservation of CPβ. The secondary structure elements of *Pb*CPβ are indicated on top of the alignment. The aligned sequences (PbCPb: A0A509AQN8, PfCPb: Q8I3T2, GgCPb: P14315, SsCPb: A9XFX6, DdCPb: P13021) were grouped and colored respective to a Risler matrix, using the *ESPript* convention. Residues important for canonical intraheterodimer contacts, actin-, CARMIL-, V1/myotrophin-, and $PIP_2$-binding are indicated by black, magenta, cyan, green, and orange dots, respectively.

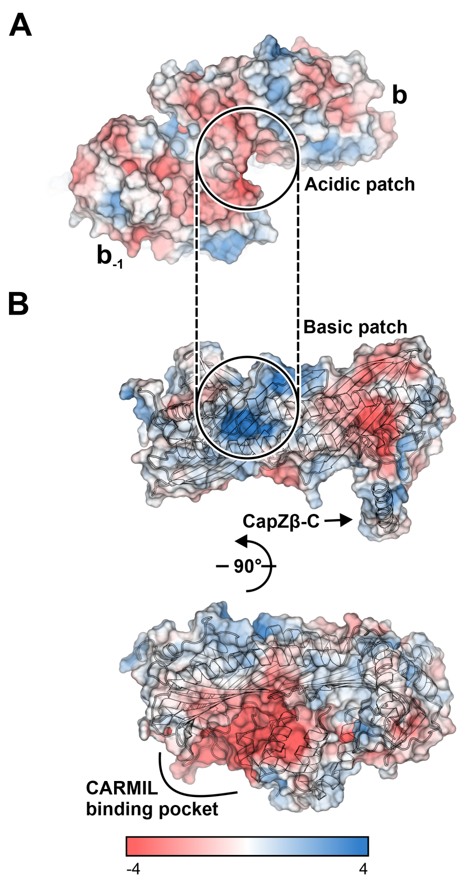


**Figure S4.** Electrostatic potential surface of canonical cytoplasmic β-actin and CapZαβ. (A) Electrostatic surface of the last (b) and penultimate ($b_{-1}$) β-actin protomer (PDB ID: 6F1T) at the barbed end. (B) Electrostatic surface of CapZαβ (PDB ID: 1IZN).

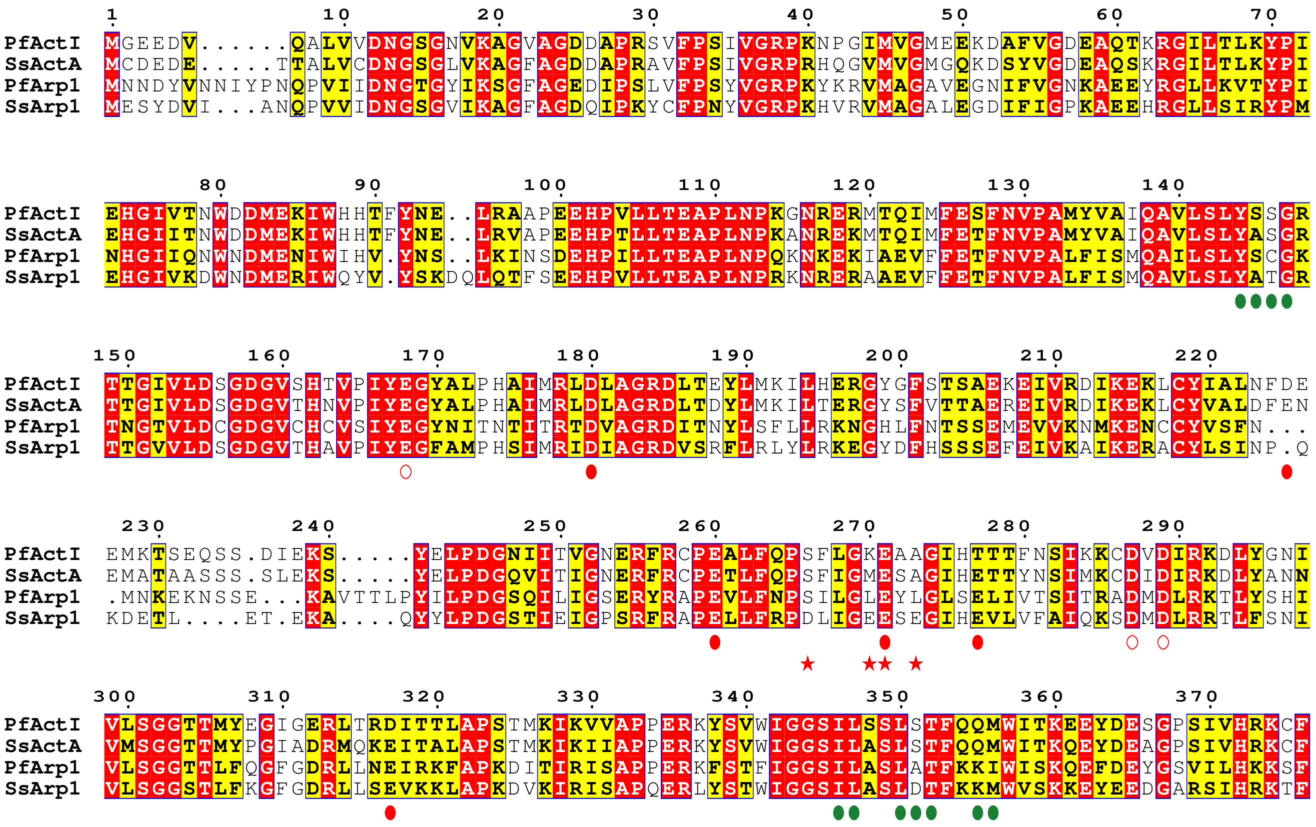


**Figure S5.** Sequence conservation of actin and Arp1. The aligned sequences (PfActI: Q8I4X0, SsActA: P68137, PfArp1: Q8I2A2, SsArp1: F2Z5G5) were grouped and colored respective to a Risler matrix, using the *ESPript* convention. Red and green symbols show residues involved in forming the acidic or hydrophobic patch, respectively. Full dots or open circles denote residues on the last or penultimate actin filament protomer, respectively. Stars mark additional acidic residues on the last Arp1 monomer in the dynactin structure (8).

**Table SI.** Surface and interface areas of the CP dimers. The letters after PDB ID denote the protein chains used in the calculations (6, 10, 11). ASA: accessible surface area, BSA: buried surface area, HB: hydrogen bonds, SB: salt bridges.

| | **1IZN_AB** | **4AKR_AB** | **6F1T_KL** | **7A0H_AB** |
|---|---|---|---|---|
| **Subunit ASA** ($Å^2$) | 17200/16900 | 15400/14700 | 16800/16600 | 17400/17200 |
| **Subunit BSA** ($Å^2$) | 3900/3700 | 3700/3500 | 3700/3600 | 2300/2400 |
| **Total ASA** ($Å^2$) | 26600 | 23000 | 26800 | 30600 |
| **Total BSA** ($Å^2$) | 7600 | 7200 | 7300 | 4700 |
| **HB/SB** | 45/6 | 42/9 | 44/10 | 18/0 |

**Table SII.** R.m.s.d. values of different CP structures superimposed on each other. **(A)** Comparison of CP dimers. **(B)** Comparison of CP subunits. **(C)** Comparison of *Pb*CPαα$^{\Delta C20}$ domains to CP subunits. The letters after the PDB IDs denote the protein chains used in the calculations (6, 10, 11). The r.m.s.d. values are given in Å and colored from green to red relative to the values in the entire table.

**A**

| CP dimers | 1IZN_AB | 4AKR_AB | 6F1T_KL | 7A0H_AB |
|---|---|---|---|---|
| **1IZN_AB** | 0.0 | 1.8 | 3.0 | 4.9 |
| **4AKR_AB** | 1.8 | 0.0 | 2.3 | 5.0 |
| **6F1T_KL** | 3.0 | 2.3 | 0.0 | 4.6 |
| **7A0H_AB** | 4.9 | 5.0 | 4.6 | 0.0 |

**B**

| CP subunits | 1IZN_A | 1IZN_B | 4AKR_A | 4AKR_B | 6F1T_K | 6F1T_L | 7A0H_A | 7A0H_B |
|---|---|---|---|---|---|---|---|---|
| **1IZN_A** | 0.0 | 4.4 | 1.7 | 3.8 | 2.3 | 4.7 | 2.9 | 3.5 |
| **1IZN_B** | 4.4 | 0.0 | 3.5 | 1.1 | 4.7 | 2.6 | 4.3 | 3.9 |
| **4AKR_A** | 1.7 | 3.5 | 0.0 | 3.4 | 2.2 | 4.5 | 2.9 | 3.3 |
| **4AKR_B** | 3.8 | 1.1 | 3.4 | 0.0 | 3.7 | 2.1 | 4.3 | 3.9 |
| **6F1T_K** | 2.3 | 4.7 | 2.2 | 3.7 | 0.0 | 4.7 | 3.0 | 3.3 |
| **6F1T_L** | 4.7 | 2.6 | 4.5 | 2.1 | 4.7 | 0.0 | 4.6 | 4.2 |
| **7A0H_A** | 2.9 | 4.3 | 2.0 | 4.3 | 3.0 | 4.6 | 0.0 | 2.8 |
| **7A0H_B** | 3.5 | 3.9 | 3.3 | 3.9 | 3.3 | 4.2 | 2.8 | 0.0 |

**C**

| CP domains | | 1IZN_A | 1IZN_B | 4AKR_A | 4AKR_B | 6F1T_K | 6F1T_L | 7A0H_A | 7A0H_B |
|---|---|---|---|---|---|---|---|---|---|
| **7A0H_A** | **Stalk** | 2.3 | 1.8 | 2.4 | 2.4 | 2.0 | 1.8 | 0.0 | 1.2 |
| | **Globule** | 2.0 | 2.8 | 2.0 | 2.8 | 2.4 | 2.9 | 0.0 | 1.4 |
| | **C. sheets** | 2.7 | 2.9 | 2.5 | 2.8 | 2.9 | 3.4 | 0.0 | 2.9 |
| | **H5 helix** | 1.3 | 2.0 | 1.2 | 1.5 | 2.1 | 2.0 | 0.0 | 1.0 |
| **7A0H_B** | **Stalk** | 2.6 | 2.0 | 2.5 | 1.8 | 2.5 | 2.0 | 1.2 | 0.0 |
| | **Globule** | 2.4 | 2.9 | 2.1 | 2.9 | 1.9 | 3.0 | 1.4 | 0.0 |
| | **C. sheets** | 2.8 | 2.8 | 2.8 | 2.5 | 2.6 | 2.7 | 2.9 | 0.0 |
| | **H5 helix** | 1.9 | 2.2 | 2.0 | 2.3 | 2.5 | 2.0 | 2.3 | 0.0 |
